# Effects of in vitro hemolysis and repeated freeze-thaw cycles in protein abundance quantification using the SomaScan and Olink assays

**DOI:** 10.1101/2024.09.21.613295

**Authors:** Julián Candia, Giovanna Fantoni, Ruin Moaddel, Francheska Delgado-Peraza, Nader Shehadeh, Toshiko Tanaka, Luigi Ferrucci

## Abstract

SomaScan and Olink are affinity-based platforms that aim to estimate the relative abundance of thousands of human proteins with a broad range of endogenous concentrations. In this study, we investigated the effects of *in vitro* hemolysis and repeated freeze-thaw cycles in protein abundance quantification across 10,776 (11K SomaScan) and 1472 (Olink Explore 1536) analytes, respectively. Using SomaScan, we found two distinct groups, each one consisting of 4% of all aptamers, affected by either hemolysis or freeze-thaw cycles. Using Olink, we found 6% of analytes affected by freeze-thaw cycles and nearly half of all measured probes significantly impacted by hemolysis. Moreover, we observed that Olink probes affected by hemolysis target proteins with a larger number of annotated protein-protein interactions. We found that Olink probes affected by hemolysis were significantly associated with the erythrocyte proteome, whereas SomaScan probes were not. Given the extent of the observed nuisance effects, we propose that unbiased, quantitative methods of evaluating hemolysis, such as the hemolysis index successfully implemented in many clinical laboratories, should be adopted in proteomics studies. We provide detailed results for each SomaScan and Olink probe in the form of extensive Supplementary Data files to be used as resources for the growing user communities of both platforms.

## Introduction

It is well known that pre-analytical variation, defined as differences in collection, handling, storage, and sample processing, can affect the measurement of protein abundances, therefore impacting the scientific discovery potential of proteomics assays. On the one hand, pre-analytical variation may manifest itself in the form of *in vitro* or spurious hemolysis.^1–4^ Indeed, estimated to range between 1 and 5% of the total samples processed in clinical settings, hemolyzed specimens are reportedly the most prevalent manifestation of procedural mistakes in laboratory testing. ^5^ On the other hand, pre-analytical variation may occur also as a result of repeated freeze-thaw cycles (FTC). Typically, blood samples are stored in −80 *^◦^*C freezers prior to analysis; in practice, however, biobanked samples may undergo multiple cycles of aliquot thawing and refreezing. Previous studies have investigated FTC effects on a variety of physiological compartments by different experimental approaches. ^6–12^ Among them, Wang et al.^6^ explored FTC effects on lipoproteins and metabolites in serum and urine samples profiled by proton nuclear magnetic resonance (NMR) and reported minor but significant effects on some of them. Mitchell et al. ^7^ investigated the impact of FTC on the plasma proteome characterized by the matrix assisted laser desorption/ionization time-of-flight (MALDI-TOF) mass spectrometry approach and observed a trend towards increasing changes in peak intensity. Fliniaux et al. ^8^ reported that 5 to 10 FTC influenced the proton NMR metabolic profile of serum samples. In this context, the aim of our study was to explore the concurrent effects of *in vitro* hemolysis and repeated FTC in protein abundance quantification using SomaScan^13^ and Olink,^14^ which are, respectively, the leading aptamer- and antibody-based platforms in current proteomics research.

SomaScan^13^ is a highly multiplexed, aptamer-based assay that targets thousands of human proteins broadly ranging from femto- to micro-molar concentrations. This technology relies on protein-capture SOMAmer (*Slow Offrate Modified Aptamer*) reagents,^15^ designed to optimize high affinity, slow off-rate, and high specificity to target proteins. These targets extensively cover major molecular functions including receptors, kinases, growth factors, and hormones, and span a diverse collection of secreted, intracellular, and extracellular proteins or domains. In this study, we utilized the 11K (v5.0) SomaScan assay, which is the most recent version of this platform, capable of measuring 10,776 different SOMAmers targeting human proteins.^16^ Olink,^14^ in turn, implements the Proximity Extension Assay (PEA) technology, a dual-recognition immunoassay in which two matched antibodies labeled with unique DNA oligonucleotides simultaneously bind to a target protein in solution. By bringing the two antibodies into proximity, their DNA oligonucleotides are allowed to hybridize, serving as a DNA barcode to identify and quantify the abundance of a specific target antigen. In this study, we utilized the Olink Explore 1536 assay to measure 1472 analytes across four panels, namely cardiometabolic, inflammation, neurology, and oncology.

Multiple studies have previously analyzed the sensitivity and specificity of SomaScan and Olink, performed direct comparisons between them, and also compared them against other proteomic platforms. SomaScan’s variability, for example, has been assessed using technical replicates in a variety of matrices (primarily plasma, serum, and cerebrospinal fluid) and assay versions spanning over a decade, from 1.1K to the most recently released 11K. The most recent study^16^ on the 11K SomaScan assay reported a median inter-plate coefficient of variation (CV) of 5%, which appears consistent with CV estimates performed in previous versions of the assay.^17–23^ Studies performed on Olink Explore 1536 (carried out in this study) and/or 3072 showed somewhat lower precision. ^24–28^ Furthermore, a growing body of literature has been devoted to comparing proteomic measurements across platforms. Emilsson et al.^29^ ran a Novartis custom-designed 5K serum SomaScan assay on 5,457 participants from the AGES Reykjavik study and performed a direct validation using two different mass spectrometry techniques, namely data dependent analysis (DDA) and multiple reaction monitoring (MRM). Results of the mass spectrometry experiments provided confirmatory evidence of 779 SOMAmer reagents binding their endogenous targets (736 by DDA and 104 by MRM). Similarly, by comparing Olink panels with DDA-MS and DIA-MS, Petrera et al.^30^ reported that the reproducibility of distributions between PEA and MS was as high as within MS platforms. Sun et al.^31^ performed 4K plasma SomaScan on 3,301 healthy participants from the INTERVAL study. They identified 1,927 genotype–protein associations (pQTLs), and tested 163 of them with Olink, finding strongly correlated effect-size estimates (*r* = 0.83). To assess potential off-target cross-reactivity, they also tested 920 SOMAmers for detection of proteins with at least 40% sequence homology to the target protein. Although 126 (14%) SOMAmers showed comparable binding with a homologous protein, nearly half of these were binding to alternative forms of the same protein. Pietzner et al.^21^ used SomaScan and Olink to create a genetically anchored cross-platform proteome-phenome network comprising 547 protein–phenotype connections, 36.3% of which were only seen with one of the two platforms suggesting that both techniques capture distinct aspects of protein biology. Their results showcased the synergistic nature of aptamer- and antibody-based proteomic profiling technologies to better understand human health and identify disease mechanisms. Other studies have performed comparisons between SomaScan, Olink, and a variety of immuno-assays, including ELISA, Myriad RBM Luminex, Meso Scale Discovery, ProterixBio, Milliplex, Randox Biochip Array, Abbott Architect, and UniCel, across multiple cohorts.^19–21,32–36^ All these platforms span a broad range in measurement precision, proteome coverage, protein target specificity, ability to uncover phenotypic associations and sensitivity to proteoforms, besides other technical considerations such as experimental setup requirements, cost of kits, reagents, and labor, and volume of sample required. Therefore, these proteomic platforms can be used complementarily with each other, as weaknesses in one of them are strengths in another. ^28,37^

To the best of our knowledge, however, no studies to date have been dedicated to investigate the impact of pre-analytical effects on protein abundance quantification measurements obtained with either platform. In this context, we explored the concurrent effects of *in vitro* hemolysis and repeated FTC in protein abundance quantification using SomaScan and Olink. To that end, we analyzed a total of 90 plasma samples from 15 healthy adult human donors evenly distributed in age and sex under different hemolysis and FTC conditions. By implementing mixed-effects models, we identified analytes showing significant effects resulting from pre-analytical perturbations. Furthermore, we mapped all combinations of SomaScan and Olink probes to unique UniProt IDs and investigated possible associations with protein characteristics obtained from the UniProt Knowledgebase. Detailed results are provided in the form of Supplementary Data files to be used by the SomaScan and Olink research communities as technical resource materials.

## Experimental Section

### Study design

Fig. 1 shows details of the study design. Blood samples were collected from 15 adult participants enrolled in the Baltimore Longitudinal Study of Aging^38^ (BLSA), a study of normative human aging established in 1958 and conducted by the National Institute on Aging, NIH. The study protocol was conducted in accordance with Declaration of Helsinki principles and was reviewed and approved by the Institutional Review Board of the NIH’s Intramural Research Program. Written informed consent was obtained from all participants. Samples used in this study were obtained from recent visits scheduled between February and April 2023. The participants’ sex and age were approximately evenly distributed, namely: 7 females (47%) with ages in the 24-89 years old range (mean: 54.6 y.o., median: 51 y.o.) and 8 males (53%) with ages in the 25-95 years old range (mean: 58.9 y.o., median: 56.5 y.o.). Each biological sample was split into six technical replicates obtained from combining the hemolyzed/non-hemolyzed condition with 3, 10 and 20 freeze-thaw cycles, which resulted in a total of 90 samples. These samples were then split into two aliquots and placed into separate 96-well plates to be analyzed in parallel with the SomaScan and Olink proteomics platforms. Because of calibrators, quality control and buffer wells, a single SomaScan plate is only able to accommodate up to 85 user-provided samples; we thus randomly selected 5 samples to be removed from the SomaScan study. Olink, on the other hand, was able to fit all 90 samples in a single plate (after removing two sample controls).

**Figure 1:**
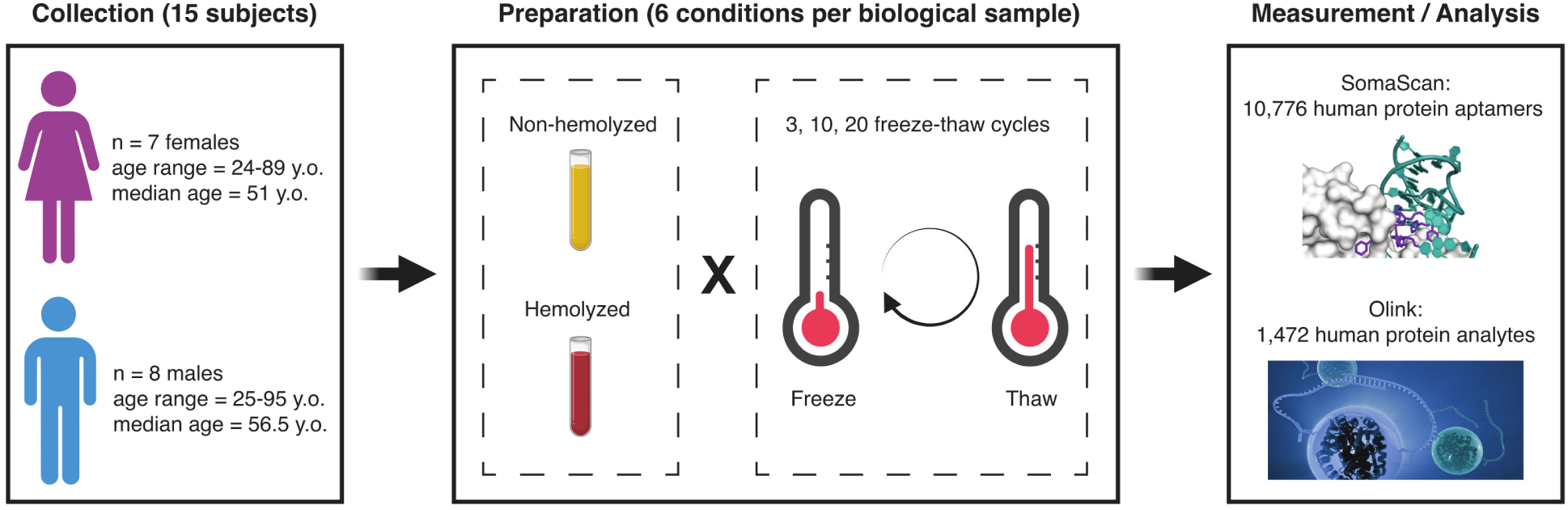
Study design. Blood samples were collected from 15 participants in the Baltimore Longitudinal Study of Aging (BLSA), approximately evenly distributed across sex and age. Each biological sample was split into six technical replicates obtained from combining the hemolyzed/non-hemolyzed condition with 3, 10 and 20 freeze-thaw cycles. The resulting samples were then analyzed with the SomaScan and Olink proteomics platforms.

### Sample preparation

From each participant, fasting whole blood was collected by venipuncture into two (3.0 mL K2 EDTA BD Vacutainer) collection tubes. Hemolysis was induced in one of the collection tubes by forcing blood through 28G needles several times until hemolysis was achieved. Samples were visually inspected to ensure they were hemolyzed (Supplementary Fig. 1) and later confirmed via Hemolysis Index (*HI*) estimates (Supplementary Data 1). The blood was centrifuged at 1200 x g at 20 *^◦^*C for 15 minutes to obtain plasma. Three 500 *µ*L aliquots of plasma were obtained from each tube and stored at −80 *^◦^*C until assayed. For FTC, each aliquot was put in dry ice for 15 minutes and subsequently left at room temperature until thawed and additional 10 minutes after complete thawing. This procedure was performed 3, 10, or 20 times depending on the condition. It should be noted that biobanked samples from long-running longitudinal studies may be stored in single, large-volume vials, which may have been thawed and re-freezed multiple times in the course of several decades. While other studies have previously explored effects of up to five^6,7^ or ten^8^ freeze-thaw cycles, our study design aimed to expand that range to encompass worst-case scenarios.

### Hemolysis Index

The Hemolysis Index (*HI*) was determined based on previous work with slight modifications.^39^ Briefly, 5 *µ*l of plasma were added to 20 *µ*l of PBS 1x for a 5-fold dilution and added onto a black, 96-well plate with a clear bottom (Corning, Inc.; Corning, NY, USA). Absorbance was measured using the Synergy H1 microplate reader (Biomek, Beckman Coulter, Inc.; Brea, CA, USA) at wavelengths of (*A*) 658 nm and (*B*) either 410 nm, 451 nm, 545 nm, or 571 nm. *HI* was calculated as *HI* = 440.54×(Absorbance(B) − 0.8967 × Absorbance(A)). The association between hemolyzed samples and *HI* was tested via mixed-effects models under different scenarios (see “Statistical methods” below) and found to be very significant (p − value *<* 10^−12^). Results are provided in Supplementary Data 1.

### The SomaScan assay

Proteins were measured using the 11K (v5.0) SomaScan assay (SomaLogic, Inc.; Boulder, CO, USA). This technology relies on aptamers (SOMAmers), which are single-stranded, chemically-modified nucleic acids, selected via the SELEX (Systematic Evolution of Ligands by EXponential enrichment) process, designed to optimize high affinity, slow off-rate, and high specificity to target proteins. The 11K assay consists of 10,776 SOMAmers targeting annotated human proteins. In order to cover a broad range of endogenous concentrations, SOMAmers are binned into different dilution groups, namely 20% (1:5) dilution for proteins typically observed in the femto- to pico-molar range (which comprise 80% of all human protein SOMAmers in the assay), 0.5% (1:200) dilution for proteins typically present in nano-molar concentrations (18% of human protein SOMAmers in the assay), and 0.005% (1:20,000) dilution for proteins in micro-molar concentrations (2% of human protein SOMAmers in the assay). The human plasma volume required is 55 *µ*L per sample. The experimental procedure follows a sequence of steps, namely: (1) SOMAmers are synthesized with a fluorophore, photocleavable linker, and biotin; (2) diluted samples are incubated with dilution-specific SOMAmers bound to streptavidin beads; (3) unbound proteins are washed away, and bound proteins are tagged with biotin; (4) UV light breaks the photocleavable linker, releasing complexes back into solution; (5) non-specific SOMAmer-protein complexes dissociate while specific complexes remain bound; (6) a polyanionic competitor is added to prevent rebinding of non-specific SOMAmer-protein complexes; (7) biotinylated proteins (and bound SOMAmers) are captured on new streptavidin beads; and (8) after SOMAmers are released from the complexes by denaturing the proteins, fluorophores are measured following hybridization to complementary sequences on a microarray chip. It should be pointed out that steps (5-6) take advantage of solution kinetics to selectively enrich for specificity, since fast off-rate, non-specific SOMAmer-protein complexes quickly dissociate, and rebinding is blocked by the polyanionic competitor. Because dissociation rates of cognate SOMAmer-protein interactions are much slower than those of non-specific interactions, specific complexes remain tightly bound on this timescale. The fluorescence intensity detected on the microarray in step (8), measured in *RFU* (*Relative Fluorescence Units*), is assumed to reflect the target epitope abundance in the original sample. In order to account for intra- and inter-plate effects, a normalization procedure is applied, consisting of a sequence of steps, whose main elements are hybridization normalization, median signal normalization, plate-scale normalization, and inter-plate calibration. ^17,22^ Results in this paper are based on the fully normalized data. For downstream analyses, protein relative concentration was assumed as given by log_10_(*RFU*). Further information about the SomaScan assay is available on the manufacturer’s website (www.somalogic.com).

### The Olink assay

Proteins were measured using Olink Explore 1536 (Olink Proteomics AB, Uppsala, Sweden) according to the manufacturer’s instructions. This assay comprises four panels focused on different disease and biological processes, namely Cardiometabolic (369 analytes), Inflammation (368 analytes), Neurology (367 analytes), and Oncology (368 analytes), for a total of 1472 analytes. For each panel, the human plasma volume required is 2.8 *µ*L per sample. The technology behind the Olink protocol is based on the Proximity Extension Assay (PEA),^14^ which is coupled with readout via next-generation sequencing. In brief, pairs of oligonucleotide-labeled antibody probes designed for each protein bind to their target, bringing the complementary oligonucleotides in close proximity and allowing for their hybridization. The addition of a DNA polymerase leads to the extension of the hybridized oligonucleotides, generating a unique protein identification “barcode”. Next, library preparation adds sample identification indices and the required nucleotides for Illumina sequencing. Prior to sequencing using the Illumina NovaSeq 6000, libraries go through a bead-based purification step and the quality is assessed using the Agilent 2100 Bioanalyzer (Agilent Technologies, Palo Alto, CA, USA). The raw count data was generated using bcl2counts (v2.2.0) and was quality controlled, normalized and converted into Normalized Protein eXpression (*NPX*), Olink’s proprietary unit of relative abundance. Data normalization is performed using an internal extension control and an external plate control to adjust for intra- and inter-run variation. All assay validation data (detection limits, intra- and inter-assay precision data, predefined values, etc.) are available on the manufacturer’s website (www.olink.com). It should be noted that, while there was an inaccuracy for the plate control on one of the four blocks on the oncology panel, the plate controls were precise based on MAD-Z score. The plate data, limits of detection, and correlation assay between Neurology and Oncology plates did not indicate issues with sample data quality.

### Protein annotations

SOMAmers are uniquely identified by their aptamer sequence ID (“SeqId”), but the relationship between SOMAmers and annotated proteins is not one-to-one. In the 11K assay, the 10,776 SOMAmers that target human proteins are mapped to 9609 unique UniProt IDs, yielding a total of 10,893 unique SOMAmer-UniProt ID pairs. Supplementary Data 2 provides the list of SOMAmer-UniProt ID pairs, along with protein annotations extracted from the UniProt Knowledgebase (UniProtKB), including length (number of amino acids in the canonical sequence), mass (molecular weight), pH dependence, Redox potential, temperature dependence, protein interactions, and subcellular location. Analogously, the 1472 Olink probes are mapped to 1465 unique UniProt IDs, yielding 1476 unique Olink-UniProt ID pairs, which are listed in Supplementary Data 3 along with UniProtKB annotations. The overlap between the 11K SomaScan and the four Olink panels used in our study comprises 1872 SOMAmers and 1353 Olink probes mapped to 1356 unique UniProt IDs, yielding a total of 1893 SOMAmer/Olink/UniProt ID triplets, listed in Supplementary Data 4. Notice that the list comprises 1891 unique SOMAmer/Olink pairs; the “Duplicated” column flags the two duplicated SOMAmer/Olink pairs. Below we describe an analysis to find SOMAmer/Olink pairs that are affected by hemolysis and freeze-thaw cycles in both platforms (see Fig. 3); results are provided in the “Paired signif.” column, indicating also the direction of change (for instance, “H: O+S-” indicates a pair that is affected by hemolysis, such that the Olink probe is increased and the paired SOMAmer decreased in hemolyzed samples). It should be noted that, out of the 1356 UniProt IDs listed in Supplementary Data 4, 410 of them are mapped to more than one SOMAmer or Olink probe, while the remaining 946 are mapped to exactly one SOMAmer and one Olink probe; UniProt IDs mapped to multiple SOMAmers and/or Olink probes are indicated by the “Multiple” column. Protein interactions obtained from the UniProt Knowledgebase are derived from known and predicted interactions, including direct (physical) and indirect (functional) associations, aggregated from multiple protein-protein interaction databases, namely: STRING, ^40^ BioGRID^41^ (Biological General Repository for Interaction Datasets), DIP^42^ (Database of Interacting Proteins), IntAct^43^ (Protein interaction database and analysis system), MINT^44^ (Molecular INTeraction database), ELM^45^ (Eukaryotic Linear Motif resource for Functional Sites in Proteins), CORUM^46^ (Comprehensive resource of mammalian protein complexes), and ComplexPortal. ^47^

### Statistical methods

For each probe in the SomaScan and Olink assays, mixed-effects models were run according to the formula:

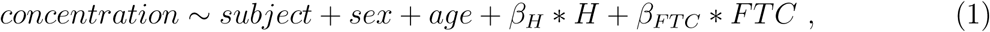

where hemolysis (*H*) and freeze-thaw cycles (*FTC*) were treated as fixed effects, and subject ID, sex and age were considered random effects. Age was discretized using tertile bins; hemolysis was coded as *H* = 1 for hemolyzed samples and *H* = 0 for non-hemolyzed samples, and the freeze-thaw cycle variable was defined as *FTC* = 1, 2, and 3 corresponding to the three conditions performed (3, 10, and 20 cycles, respectively). Mixed-effects models were implemented in a script via the lmer function from R package lme4 v.1.1.35.5 with the argument REML=FALSE to optimize the log-likelihood.^48^ The statistical significance of effects associated to hemolysis was assessed by performing a likelihood ratio test to compare the full model from Eq. (1) against the null model:

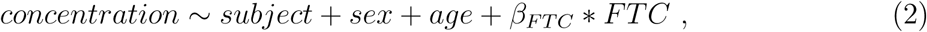

using the anova function from R package stats v.4.4.0. Similarly, the significance of effects associated with freeze-thaw cycles was derived from comparing the full model against the null model:

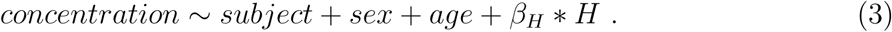

In order to check the association between hemolyzed samples and quantitative measures via the hemolysis index (*HI*), we followed a similar approach, where (i) a mixed-effects model of the form *HI* ∼ *subject* + *β_H_* ∗ *H* was compared against the null model *HI* ∼ *subject*, and (ii) a mixed-effects model of the form *HI* ∼ *subject* + *β_H_* ∗ *H* + *β_F T C_* ∗ *FTC* was compared against the null model *HI* ∼ *subject* + *β_F T C_* ∗ *FTC*. These procedures were run independently for four different measures of *HI* at different wavelengths.

### Data and Software Availability

Data analysis was performed in R v.4.4.0. Protein annotations from UniProtKB were extracted using the R package queryup v.1.0.5. Mixed-effects models were implemented via lme4 v.1.1.35.5. Following pre-processing with fgsea v.1.30.0 to import pathway gmt files and harmonize gene symbols, Gene Set Enrichment Analysis^49^ was implemented with the command-line client gsea-cli.sh v.4.3.3 and the Molecular Signatures Database hallmark gene set collection.^50^ Plots were generated using R packages RColorBrewer v.1.1-3, calibrate v.1.7.7, VennDiagram v.1.7.3, and gplots v.3.1.3.1. BioRender was used to prepare Fig. 1. Anonymized datasets and custom R source code used in our analyses are available on the Open Science Framework repository at osf.io/9rwsq (DOI 10.17605/OSF.IO/9RWSQ).

## Results and Discussion

In this study, we investigated the effects of *in vitro* hemolysis and repeated freeze-thaw cycles (FTC) in protein abundance quantification across 10,776 (SomaScan) and 1472 (Olink) analytes, respectively. Fig. 2 displays Volcano plots showing significantly affected probes, obtained from mixed-effects models separately run for SomaScan (top panels, a-b) and Olink (bottom panels, c-d), as indicated. Left-side panels (a,c) show the effects of hemolysis, while right-side panels (b,d) show the effects of repeated FTC. Detailed results are provided for SomaScan (Supplementary Data 5) and Olink (Supplementary Data 6), respectively. For SomaScan, we found 387 (4%) SOMAmers with significant effects associated to hemolysis (Fig. 2(a)), approximately evenly distributed in the positive (210 SOMAmers) and negative (177 SOMAmers) directions. A similar picture emerged when considering FTC effects: we found 466 (4%) SOMAmers affected (Fig. 2(b)), although more biased towards the negative direction (301 SOMAmers) relative to the positive one (165 SOMAmers). For Olink, in contrast, we found nearly half of the probes significantly affected by hemolysis (Fig. 2(c)). Indeed, we observed 718 (49%) affected analytes, most of which appeared to increase in hemolyzed samples (624 in the positive direction vs 94 in the negative direction). The effects of FTC (Fig. 2(d)) were much less dramatic and more in line with the observations from SomaScan; we found 84 (6%) analytes affected and biased in a similar ∼ 2 : 1 ratio towards the negative direction (58 analytes) relative to the positive one (26 analytes). We observe that PPME1 (protein phosphatase methylesterase 1) is the protein with strongest hemolysis effects in the Olink platform (*β_H_* = 4.3, p − value = 1.2 × 10^−62^) and it also displays mild but significant FTC effects (*β_F T C_* = 0.13, p − value = 0.013), both in the positive direction (i.e. increasing protein concentration with both hemolysis and FTC). On the other hand, KLK13 (kallikrein related peptidase 13) is the protein with strongest FTC effects (*β_F T C_* = −0.14, p − value = 5.4 × 10^−9^). KLK13’s concentration decreases with increasing FTC and shows no significant associations with hemolysis (p − value = 0.71). Both PPME1 (OID21347) and KLK13 (OID21291) belong to the oncology panel, an observation that may suggest that pre-analytical variation effects may be associated with the specific panels Olink probes belong to. Table 1 shows the number of probes in each panel affected by hemolysis and FTC; we did indeed observe a significant association in the former (Fisher’s exact test p − value = 4.4 × 10^−5^), showing an enrichment of affected probes in the oncology and neurology panels, whereas FTC effects did not appear to be significantly associated with panel of origin (p − value = 0.72). In a recent human plasma biomarker study^51^ using the Olink Explore 3072 platform, it was reported that 1112 probes were found to be significantly associated with hemolysis. Of those, 560 were measured in our study and 502 of them (90%) were also found to be significantly associated with hemolysis, resulting in a strong agreement between the two studies with odds ratio 27.8 (95% CI: 20.2-38.7) and Fisher’s exact test p − value *<* 2.2 × 10^−16^.

**Figure 2:**
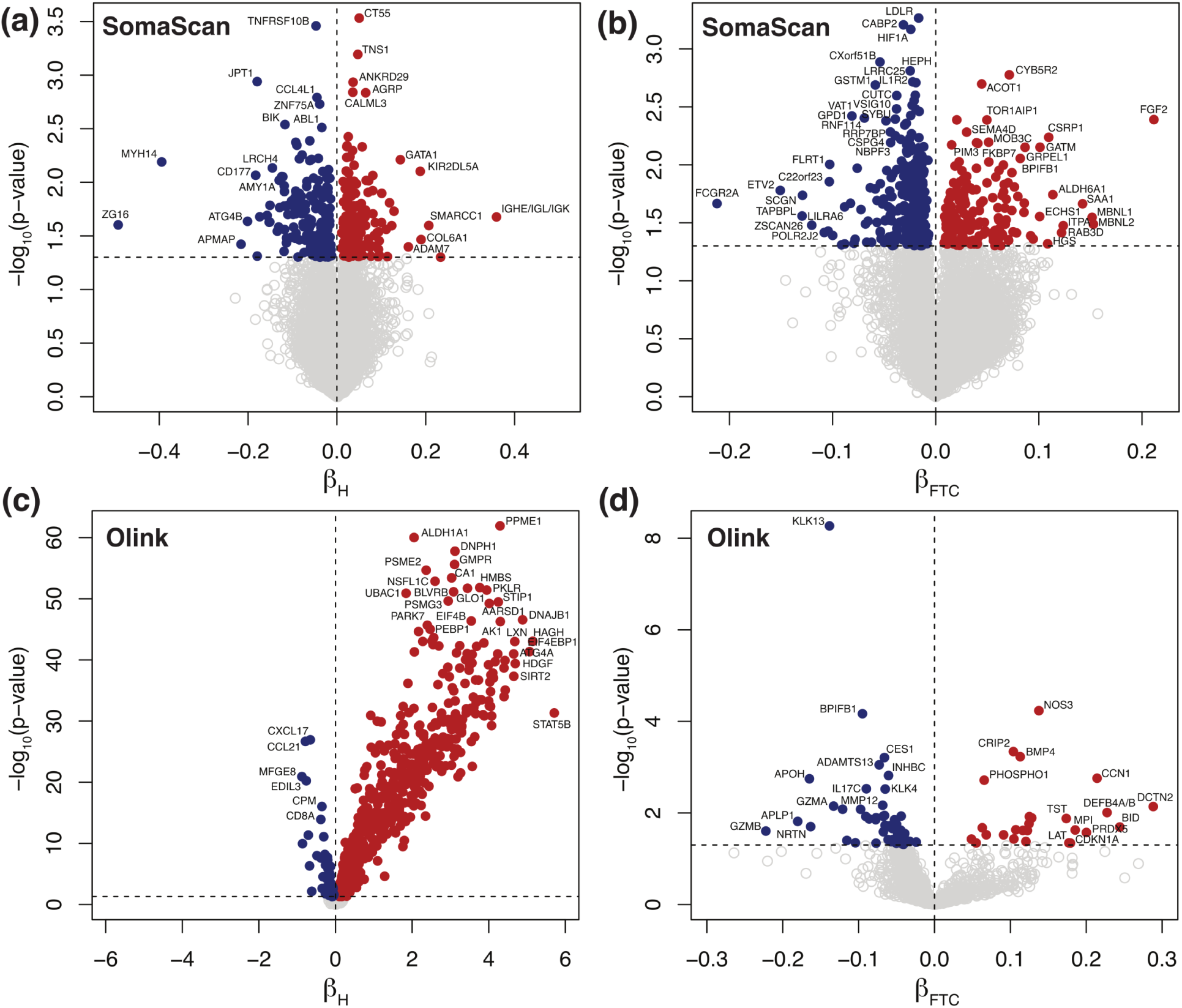
Volcano plots showing significantly affected probes. Mixed-effects models were separately run for SomaScan **(a-b)** and Olink **(c-d)**, as indicated, where symbols represent 10,776 SOMAmers and 1472 Olink probes, respectively. Left-side panels **(a,c)** show the effects of hemolysis, while right-side panels **(b,d)** show the effects of repeated freeze-thaw cycles. Significance is shown on the y-axis as unadjusted − log_10_(p − value). The horizontal dashed lines correspond to the p − value = 0.05 threshold. Probes above the significance threshold are shown as solid circles (red in the positive direction, blue in the negative direction), while those below the significance threshold are shown as open circles in light grey.

**Table 1:** Olink probes significantly affected by hemolysis and freeze-thaw cycles.

|  | **Hemolysis** |  | **Freeze-thaw cycles** |  |
| --- | --- | --- | --- | --- |
| **Panel** | **n.s.** | **significant** | **n.s.** | **significant** |
| Cardiometabolic | 225 (61%) | 144 (39%) | 349 (95%) | 20 (5%) |
| Inflammation | 192 (52%) | 176 (48%) | 350 (95%) | 18 (5%) |
| Neurology | 170 (46%) | 197 (54%) | 342 (93%) | 25 (7%) |
| Oncology | 167 (45%) | 201 (55%) | 347 (94%) | 21 (6%) |

Fig. 3 shows Venn diagrams displaying the number of probes significantly affected by hemolysis and FTC. For Olink (Fig. 3(a)), 42 (2.8%) probes are affected by both, which is a non-significant overlap (Fisher’s exact test p − value = 0.82). For SomaScan (Fig. 3(b)), only 14 (0.13%) SOMAmers are affected by both, which is also a non-significant overlap (Fisher’s exact test p − value = 0.61). In order to compare across platforms, we first needed to map Olink probes and SOMAmers to UniProt targets, then consider those UniProt targets in common to associate Olink probes to SOMAmers. This procedure yielded a total of 1893 SOMAmer/Olink/UniProt ID triplets and 1891 unique SOMAmer/Olink ID pairs, listed in Supplementary Data 4. Fig. 3(c) shows the number of affected SOMAmer/Olink ID pairs mapped via common UniProt ID targets. We found only 30 SOMAmer/Olink pairs affected by hemolysis in both platforms; of these, only 19 agree in the direction of change (17 increase, 2 decrease abundance in hemolyzed samples), whereas the remaining 11 change in opposite directions. For FTC, we only found 5 SOMAmer/Olink pairs affected, and only one of them agrees in the direction of change. Restricting the comparison to the 946 UniProt IDs mapped to exactly one SOMAmer and one Olink probe, we found 16 SOMAmer/Olink pairs affected by hemolysis in both platforms and just 2 pairs affected by FTC (Fig. 3(d)).

**Figure 3:**
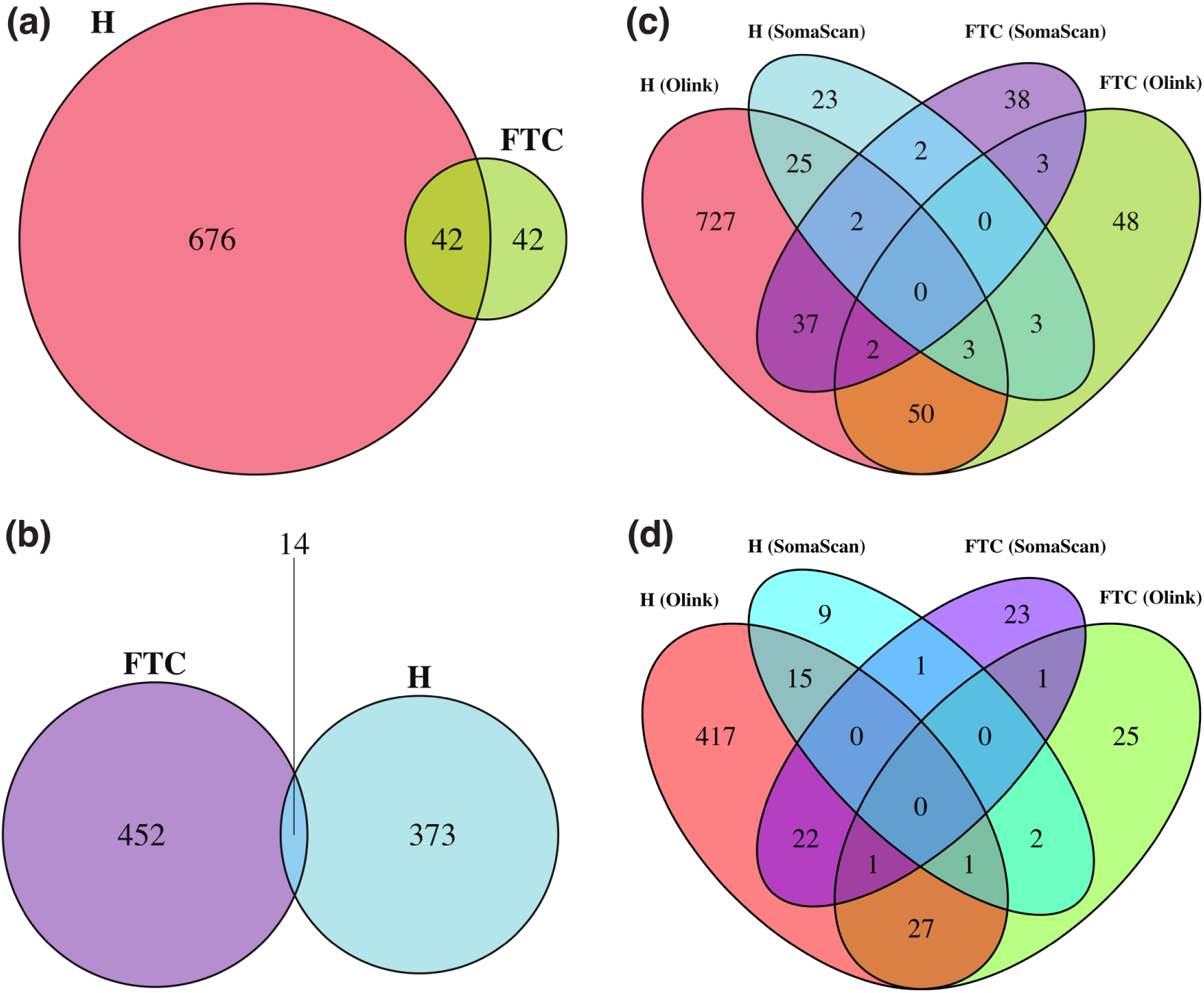
Venn diagrams showing probes significantly affected by hemolysis (H) and freeze-thaw cycles (FTC). **(a)** Number of affected Olink probes. **(b)** Number of affected SomaScan probes (SOMAmers). **(c)** Number of affected SOMAmer/Olink ID pairs mapped via common UniProt ID targets (using the full list of 1356 UniProt IDs; Supplementary Data 4). **(d)** Number of affected SOMAmer/Olink ID pairs mapped via common UniProt ID targets (using a restricted list of 946 UniProt IDs mapped exactly to just one SOMAmer and one Olink probe).

Given the large proportion of Olink probes affected by hemolysis, we investigated possible associations with protein characteristics obtained from UniProtKB (see Supplementary Table 2). We found that the number of protein interactions is larger for protein probes affected by hemolysis compared with protein probes not significantly affected by hemolysis. Table 2 shows robust summary statistics calculated over all 1476 Olink/UniProt ID pairs. The median number of interacting proteins is 2 for non-affected Olink probes and 5 for non-affected probes, while the median absolute deviation (MAD) is 2.97 and 5.93, respectively. Given this variability, we performed a Wilcoxon test (also known as Mann-Whitney U test) to assess the statistical significance of the difference in the number of interacting proteins between the two groups (or, more precisely, to test the null hypothesis that the data from the two groups originate from the same distribution). We obtained p − value = 3.5 × 10^−18^ and reject the null hypothesis; therefore, we conclude that Olink probes affected by hemolysis are associated with proteins with a larger number of annotated protein-protein interactions compared with that of Olink probes not significantly affected by hemolysis. Because a large fraction of proteins have no documented protein-protein interactions (possibly due to limitations in the available annotation databases), we repeated the analysis after removing them. Table 2 shows that the median number of interacting proteins equals 4 for Olink probes not significantly affected by hemolysis, while it equals 6 for affected probes. The Wilcoxon test yields p − value = 1.2 × 10^−11^, thus confirming the observed association. Fig. 4(a) shows the percentile distribution comparison of the number of interacting proteins for Olink targets not significantly affected by hemolysis (blue) versus those significantly affected (red). While the association between hemolysis-affected proteins and the number of interacting proteins is reported here as an exploratory result, investigating possible causal mechanisms for this observation lies beyond the scope of this paper.

**Figure 4:**
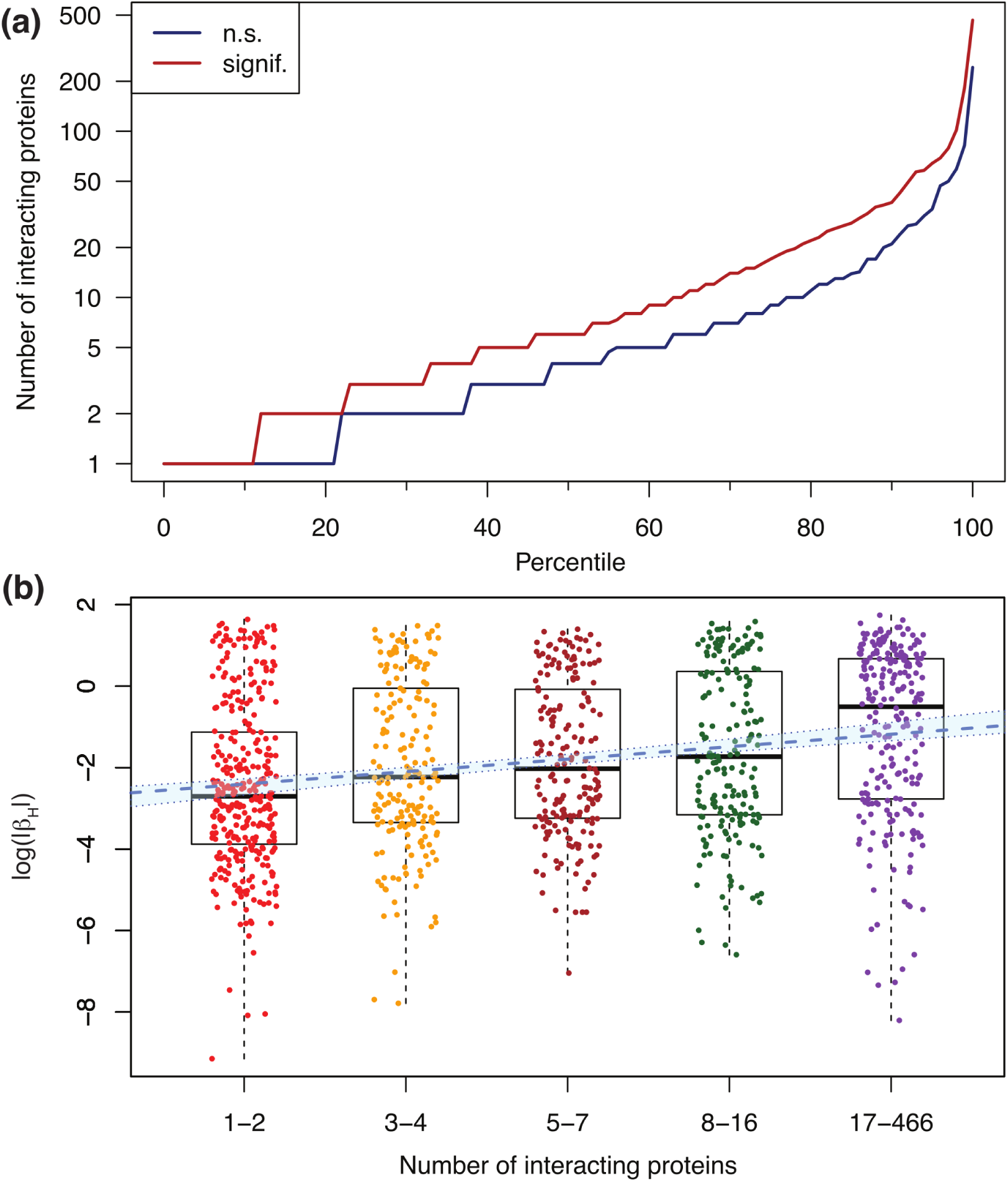
Olink probes affected by hemolysis have a larger number of annotated protein-protein interactions. **(a)** Percentile distribution comparison of the number of interacting proteins for Olink targets not significantly (n.s.) affected by hemolysis (blue) and those significantly affected by hemolysis (red). **(b)** Association between log(|*β_H_* |) and the number of interacting proteins binned in five groups of similar size. Here, *β_H_* represents the beta coefficient obtained from mixed-effects models, which captures the effect that hemolysis has on Olink’s Normalized Protein eXpression (*NPX*) metric of abundance. The blue dashed line shows the linear regression (slope=0.31, p − value = 5.9 × 10^−14^); the light-blue background enclosed within dotted boundaries shows the 95% confidence interval.

**Table 2:**
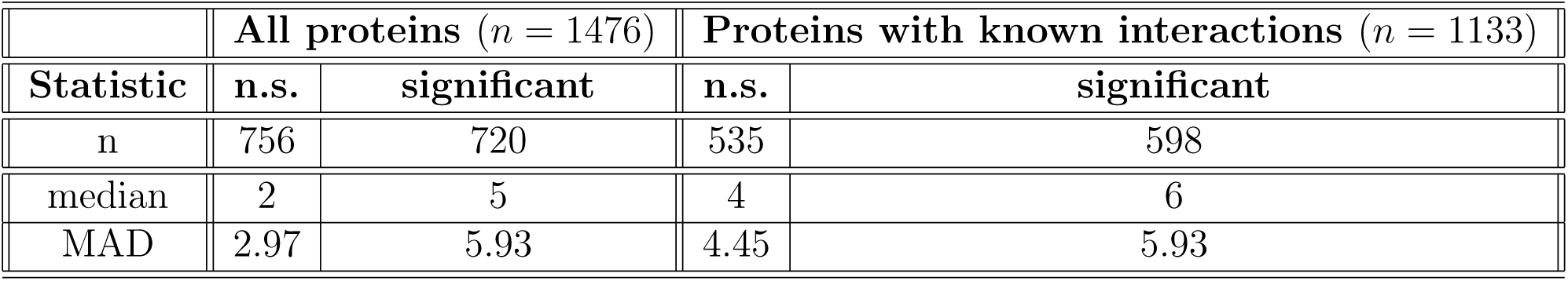
Robust summary statistics for the number of interacting proteins associated with Olink probes not-significantly (n.s.) vs significantly affected by hemolysis.

| | All proteins ($n = 1476$) | | Proteins with known interactions ($n = 1133$) | |
| --- | --- | --- | --- | --- |
| Statistic | n.s. | significant | n.s. | significant |
| n | 756 | 720 | 535 | 598 |
| median | 2 | 5 | 4 | 6 |
| MAD | 2.97 | 5.93 | 4.45 | 5.93 |

As a complementary analysis, we explored the association between the number of interacting proteins and the absolute value of beta coefficients (|*β_H_* |) obtained from the mixed effects models. We found Spearman’s correlation to be significant (*r* = 0.23 and p − value = 7.1 × 10^−15^), with the caveat that this method cannot compute exact p-values with ties. Alternatively, taking logs to account for the fact that both variables have longtailed distributions, Pearson’s correlation was found to be similarly significant (*r* = 0.22 and p − value = 4.6 × 10^−14^). Finally, by binning the number of interacting proteins among *n_b_* = 5 bins of similar size, we regressed log(|*β_H_* |) against the bin index *i_b_* = 1*, . . ., n_b_* and found a positive, significant regression slope (p − value = 5.9 × 10^−14^). The binned data with the overlaid linear fit is shown in Fig. 4(b). Hemolysis-sensitive SOMAmers, in contrast, were not found to be associated with the number of interacting proteins.

Because hemolysis is directly associated with damage to red blood cells (RBCs), we investigated the association between affected probes and the RBC proteome. ^52^ The proportion of RBC protein targets is approximately the same in both proteomics platforms; we found 2083 (19.3%) SOMAmers and 263 (17.9%) Olink probes associated with RBC proteins, which are indicated in Supplementary Data 5 and 6, respectively. Fig. 5(a) shows the contingency matrix for SOMAmers mapped to RBC proteins (rows) vs SOMAmers found to be significantly affected by hemolysis (columns). The corresponding odds ratio is 0.99 (95% CI: 0.75-1.28) with Fisher’s exact test p − value = 0.95, indicating lack of significant association for SomaScan. In contrast, results for Olink shown by Fig. 5(b) indicate a very strong association with odds ratio 8.89 (95% CI: 6.13-13.20) and Fisher’s exact test p − value *<* 2.2 × 10^−16^. Not surprisingly, the correlation of hemolysis effects between SomaScan and Olink is not significant neither across the 305 SomaScan / Olink probe pairs mapped to RBC proteins (Supplementary Fig. 2(a)) nor the subset of 199 pairs after excluding UniProt IDs that map to more than one SOMAmer or Olink probe (Supplementary Fig. 2(b)). To rule out artifacts arising from the normalization procedures of SomaScan data, we compared SOMAmer rank metrics derived from hemolysis effects using different levels of normalization and observed a strong correlation (Supplementary Fig. 3). Furthermore, Supplementary Tables 1 and 2 show hemolysis effects for probes associated with the two most abundant RBC proteins, hemoglobin and carbonic anhydrase, for the Olink and SomaScan assays, respectively. We find that Olink probes are significantly increased by hemolysis; however, that is not the case for SomaScan probes.

**Figure 5:**
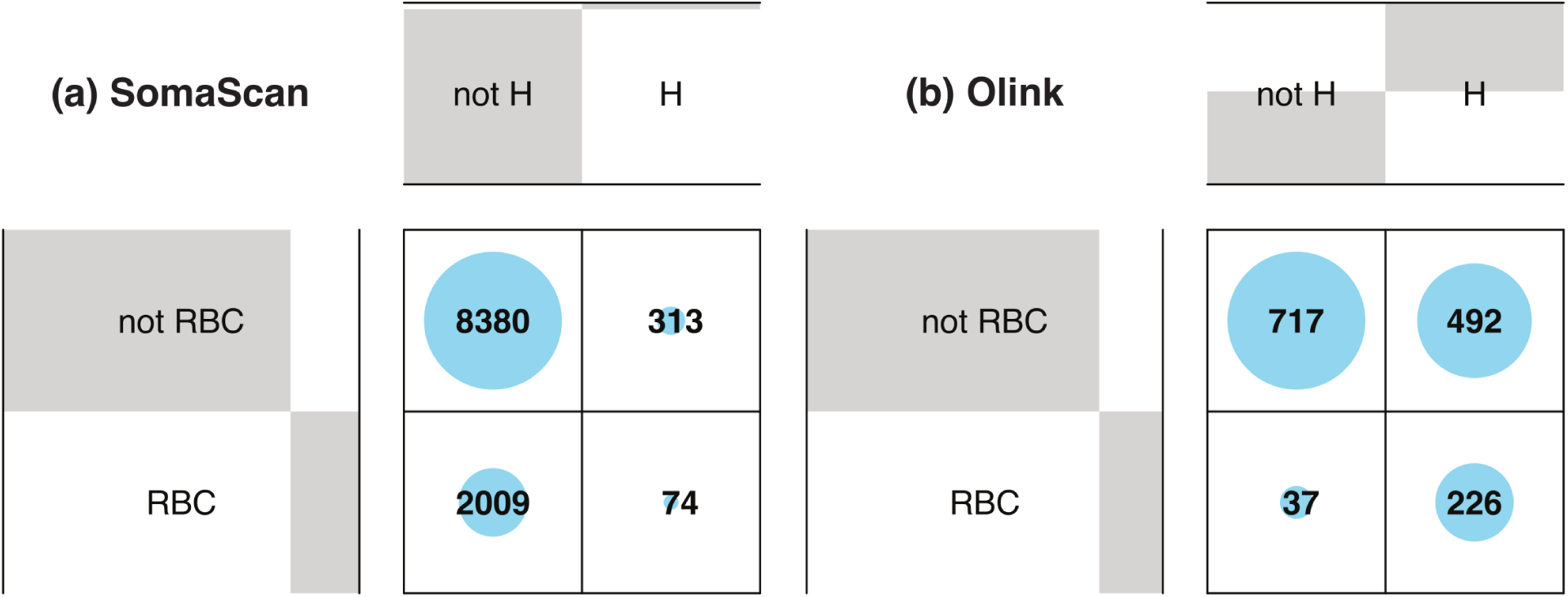
Balloon plots showing the contingency matrix between probes associated with RBC proteins (rows) and probes significantly affected by hemolysis (columns). **(a)** For SomaScan, the association is not significant: odds ratio 0.99 (95% CI: 0.75−1.28) with Fisher’s exact test p − value = 0.95. **(b)** For Olink, the association is significant: odds ratio 8.89 (95% CI: 6.13 − 13.20) and Fisher’s exact test p − value *<* 2.2 × 10^−16^.

In order to functionally characterize the proteins whose associated probes were found to be affected by hemolysis, we performed Gene Set Enrichment Analysis (GSEA)^49^ using the Hallmarks pathways^50^ as reference. Fig. 6 shows three Hallmarks pathways significantly enriched among Olink probes affected by hemolysis. The most enriched pathway comprises genes involved in the metabolism of heme (a cofactor consisting of iron and porphyrin) and erythroblast differentiation and, reassuringly, the top “core enrichment” gene in this pathway is Carbonic Anhydrase 1 (CA1), which encodes a cytosolic protein found at the highest level in erythrocytes and, as mentioned above, ranks amongst the most abundant RBC proteins. Detailed GSEA results for Olink and SomaScan probes affected by hemolysis and FTC are provided in Supplementary Data 7.

**Figure 6:**
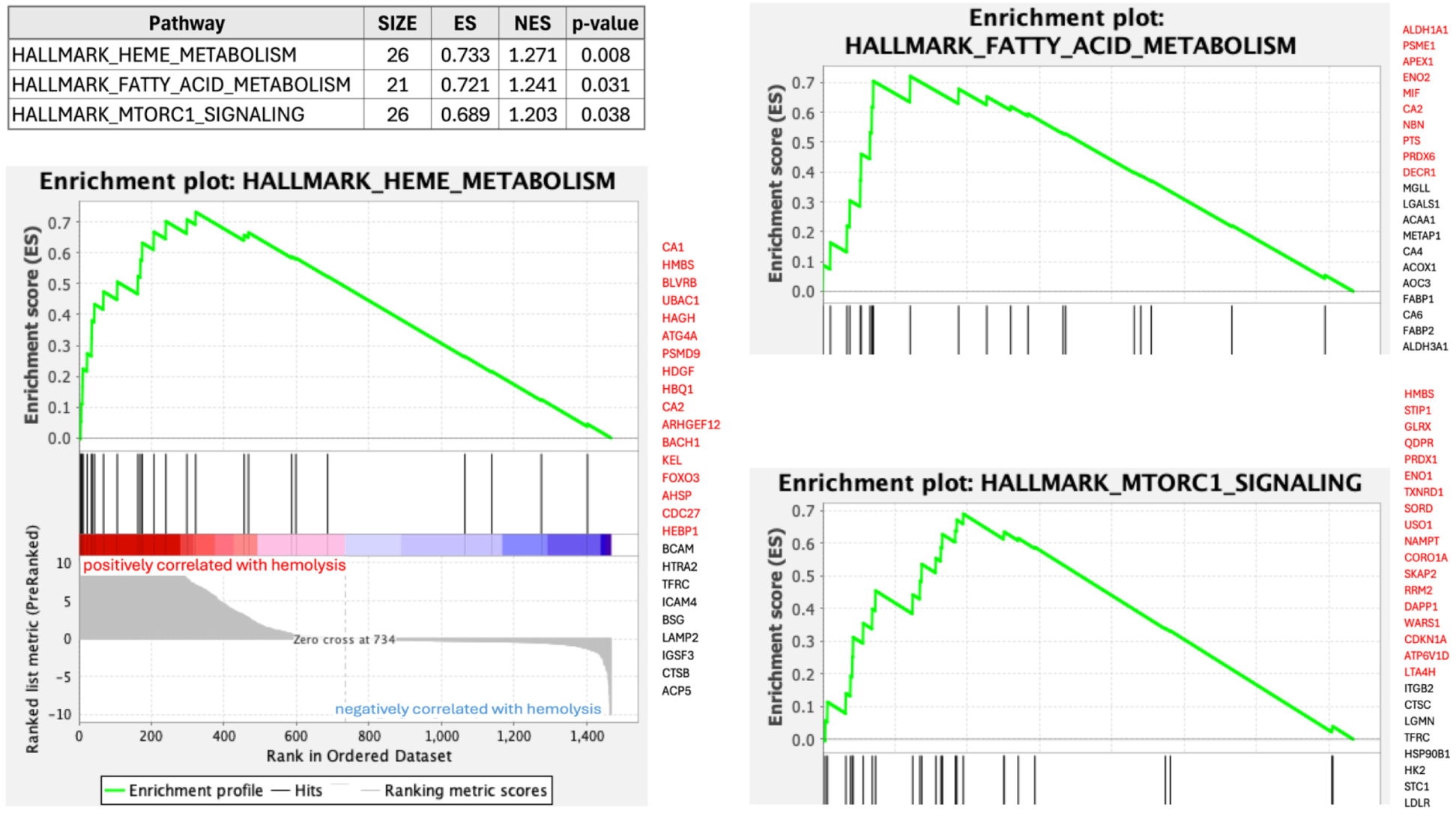
Gene Set Enrichment Analysis of Olink probes affected by hemolysis. Three Hallmarks pathways were significantly enriched. To the right hand side of each enrichment plot, we display the list of measured proteins (represented by Entrez gene symbols) in each pathway, with core enrichment genes shown in red.

Altogether, our results indicate that a large portion of the Olink probes detected at increased levels in hemolyzed samples are associated with the erythrocyte proteome.^52^ However, it should be pointed out that, although hemolyzed samples are defined by red blood cell damage, *in vitro* hemolysis is also associated with other laboratory artifacts, including damage to white blood cells, as manifested by the release of cell-free DNA^53^ and decreased white blood cells counts,^54^ which may explain the increased signals observed also in proteins not associated with RBCs. It should also be noticed that for some probes, most notably in the case of SomaScan, hemolysis has the counter-intuitive effect of decreasing the measured signal. A possible explanation for this effect is that, by the release of proteases or enzymes that add post-translational modifications to the target proteins, these may undergo structural changes that impact the affinity strength and/or the dissociation rate of the assay’s capture process. This important topic, however, should be investigated further and is out of the scope of the present paper.

## Conclusions

We investigated the effects of *in vitro* hemolysis and repeated freeze-thaw cycles in protein abundance quantification using the SomaScan and Olink assays, the leading aptamer- and antibody-based platforms in current proteomics research, respectively. For SomaScan, we found two distinct groups, each one consisting of 4% of all measured SOMAmers, affected by either hemolysis or freeze-thaw cycles. For Olink, probes affected by freeze-thaw cycles were in line with the observations from SomaScan, with 6% of probes affected and biased in a similar ∼ 2 : 1 ratio towards the negative direction (i.e. decreasing abundance by increasing freeze-thaw cycles) relative to the positive one. In contrast, however, we found that nearly half of the Olink probes were significantly affected by hemolysis and most of them biased towards the positive direction (i.e. increasing abundance by hemolysis), whereas the much smaller proportion of affected SOMAmers appeared approximately evenly distributed in both directions. Due to the large proportion of Olink probes affected by hemolysis, we investigated possible associations with protein characteristics obtained from UniProtKB and found a significant association between hemolysis-sensitive Olink probes and the number of annotated protein-protein interactions of their targets. About a third of hemolysis-affected Olink probes were associated with the erythrocyte proteome; in contrast, however, most SomaScan probes affected by hemolysis were not. Although we may conjecture that the observed differences between SomaScan and Olink may reflect differences in the detection mechanisms at play at the molecular scale between these two complementary technologies (namely, modified DNA aptamer-based vs antibody-based), this is a topic that deserves further investigation and lies beyond the scope of the present work.

Our observations highlight the need to adopt an extremely careful approach for sample pre-processing and handling, in order to ensure the integrity of proteomics measurements and downstream analysis. In clinical practice, the phenomenon of *in vitro* hemolysis affecting clinical samples is well known. Historically, hemolysis was detected by visual inspection of the specimen after centrifugation, but a newer approach is based on the hemolysis index (*HI*).^55,56^ Most clinical chemistry platforms are nowadays equipped with automated *HI* detection systems, which involve the rapid spectrophotometric measurement of free hemoglobin in serum or plasma along with decision rules for the systematic handling of specimens based on tolerance range values.^39^ We suggest that, upon further investigation, *HI*-based scores integrated with assay- and probe-specific suitable thresholds may provide an unbiased, quantitative procedure to flag and, even, correct proteomics measurements that may be affected by hemolysis. To a lesser extent, quantitative metrics to assess freeze-thaw cycles may be also implemented for similar purposes. In this regard, SomaScan has recently begun to explore and implement post-hoc, machine-learning-based estimates of pre-analytical variation called SomaSignal Tests, which aim at quantifying a variety of sample processing factors, including fed-fasted time, number of FTC, time-to-decant, time-to-spin, and time-to-freeze.^57^ While stochastic nuisance effects would theoretically even out over large datasets, it is important to consider that: (i) it is impossible to know in advance which and how many samples may potentially be affected, e.g. if a faulty protocol is implemented affecting the collection, pre-processing, and handling of the samples; (ii) the high cost of Olink and SomaScan imposes severe limits on the number of samples assayed in a single study, typically limited to one or a few plates; and (iii) the number of assayed samples may be limited by clinical study enrollment, particularly in the case of rare conditions. Therefore, the impact of potentially compromised samples due to pre-analytical variation is very high in current proteomics studies using these state-of-the-art technologies.

A limitation of this study is that we used a visual inspection approach to ensure that samples were hemolyzed (Supplementary Fig. 1). However, we did determine the *HI* after the proteomic analysis and demonstrated that there was a significant difference between hemolyzed and non-hemolyzed samples (Supplementary Data 1). We suggest that, in future work, hemolysis can be induced in a controlled manner to determine the effects of different degrees of hemolysis. Another limitation of this work is the small sample size (90 experimental samples obtained from 15 biological samples) and that, as a result, this is only an exploratory study.

Accompanying this paper, detailed results for each Olink and SOMAmer probe are provided in the form of extensive Supplementary Data files. As SomaScan and Olink become more widely adopted and utilized as state-of-the-art tools for proteomics discovery, we hope that our work will serve as a valuable technical reference and resource for the growing user communities of both platforms.

## Conflict of Interest

J.C. and L.F. have given unpaid seminars and/or webinars sponsored or co-sponsored by SomaLogic. R.M. has given unpaid seminars sponsored by Olink.

## Acknowledgement

This research was supported entirely by the Intramural Research Program of the National Institute on Aging, NIH. The authors thank Carlos Nogueras-Ortiz for useful comments on earlier versions of the manuscript draft, and two anonymous reviewers for useful and constructive criticism that helped us improve this paper.

## Supporting Information Available

The following files are available online:

- Supplementary Figure 1: 96-well plate design showing the appearance and distribution of samples.
- Supplementary Figure 2: Comparison of hemolysis effects between SomaScan and Olink probes mapped to shared RBC proteins.
- Supplementary Figure 3: Comparison of SOMAmer rank metrics of hemolysis effects using different levels of data normalization.
- Supplementary Table 1: Hemolysis effects of Olink probes mapped to hemoglobin and carbonic anhydrase proteins.
- Supplementary Table 2: Hemolysis effects of SomaScan probes mapped to hemoglobin and carbonic anhydrase proteins.
- Supplementary Data 1: Hemolysis Index (HI) measurements and mixed-effects model results to assess HI differences in hemolyzed vs non-hemolyzed samples.
- Supplementary Data 2: List of 10,893 unique SOMAmer-UniProt ID pairs, along with protein annotations extracted from UniProtKB.
- Supplementary Data 3: List of 1476 unique Olink-UniProt ID pairs, along with protein annotations extracted from UniProtKB.
- Supplementary Data 4: List of 1893 unique SOMAmer-Olink-UniProt ID triplets, along with protein data extracted from UniProtKB.
- Supplementary Data 5: Mixed-effects model results to assess the effect of hemolysis and freeze-thaw cycles on SomaScan’s protein concentration measurements.
- Supplementary Data 6: Mixed-effects model results to assess the effect of hemolysis and freeze-thaw cycles on Olink’s protein concentration measurements.
- Supplementary Data 7: Gene Set Enrichment Analysis for Olink and SomaScan probes affected by hemolysis and freeze-thaw cycles.

---

**For Table of Contents Only**

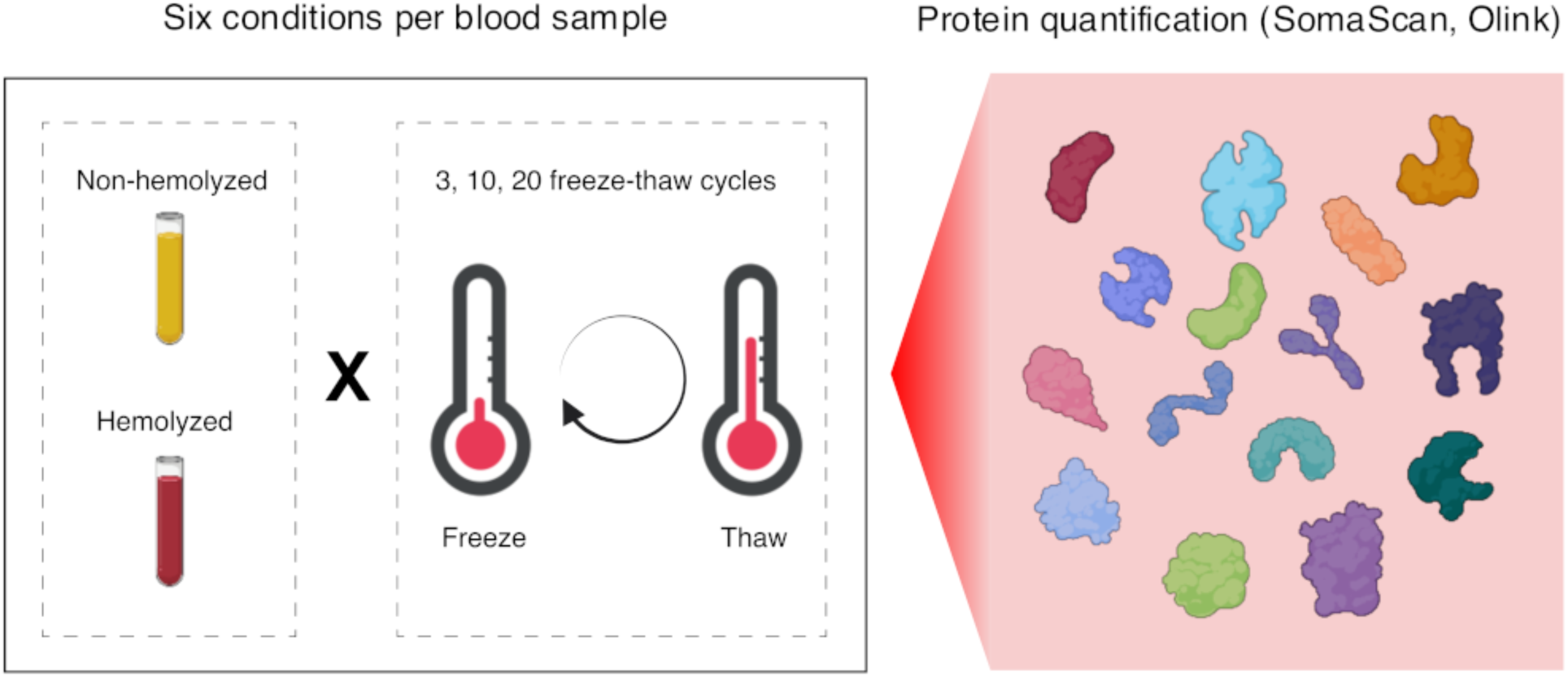

## Notes

### Summary of Updates

Minor revision: (1) Keywords were added following the Abstract section; (2) Data and software were made publicly available, adding reference to the OSF public repository; (3) References were updated, adding date of access for pre-prints.

