## Supplementary Figure 1 for "Effects of in vitro hemolysis and repeated freeze-thaw cycles in protein abundance quantification using the SomaScan and Olink assays"

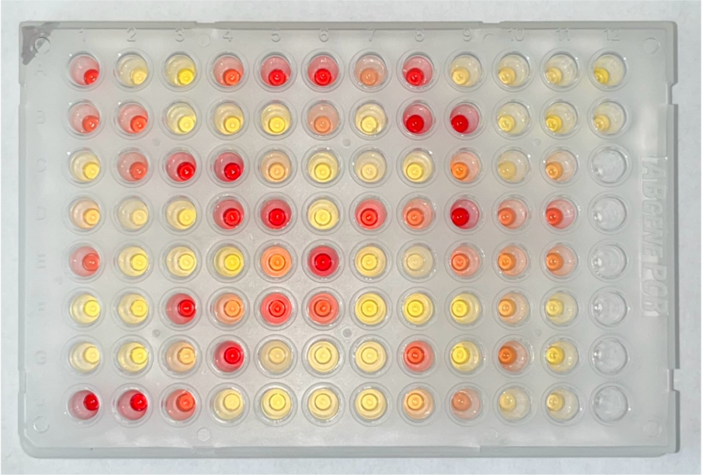

| | 1 | 2 | 3 | 4 | 5 | 6 | 7 | 8 | 9 | 10 | 11 | 12 |
|---|---|---|---|---|---|---|---|---|---|---|---|---|
| **A** | s10_10_H | s12_20_nH | s04_3_nH | s08_3_H | s07_20_H | s14_10_H | s02_10_H | s07_10_H | s15_20_nH | s15_3_nH | s07_20_nH | s04_10_nH |
| **B** | s06_3_H | s15_3_H | s13_3_nH | s01_3_nH | s08_3_nH | s02_3_H | s10_20_nH | s09_20_H | s09_3_H | s13_10_nH | s14_3_nH | s07_3_nH |
| **C** | s01_20_nH | s06_10_H | s14_3_H | s09_10_H | s13_20_H | s11_20_nH | s06_3_nH | s08_20_nH | s04_10_H | s06_10_nH | s11_20_H | |
| **D** | s13_3_H | s02_20_nH | s12_3_nH | s07_3_H | s05_10_H | s09_3_nH | s10_20_H | s12_10_H | s03_20_H | s12_20_H | s15_10_H | |
| **E** | s06_20_H | s09_10_nH | s10_10_nH | s04_20_nH | s04_3_H | s03_10_H | s01_10_nH | s06_20_nH | s12_3_H | s01_3_H | s01_20_H | |
| **F** | s11_3_nH | s08_10_nH | s14_20_H | s01_10_H | s10_3_H | s08_20_H | s05_20_nH | s05_3_nH | s14_20_nH | s11_10_H | s13_20_nH | |
| **G** | s03_10_nH | s11_10_nH | s13_10_H | s05_20_H | s02_10_nH | s07_10_nH | s15_10_nH | s08_10_H | s05_10_nH | s04_20_H | s14_10_nH | |
| **H** | s03_3_H | s05_3_H | s15_20_H | s12_10_nH | s09_20_nH | s03_3_nH | s10_3_nH | s11_3_H | s02_20_H | s02_3_nH | s03_20_nH | |

**Supplementary Figure 1. 96-well plate design showing the appearance and distribution of samples.** Samples are labeled by subject ID followed by the number of freeze-thaw cycles (3, 10, or 20) and a suffix indicating whether the sample was hemolyzed (H) or not (nH).
