## Supplementary Figure 2 for "Effects of in vitro hemolysis and repeated freeze-thaw cycles in protein abundance quantification using the SomaScan and Olink assays"

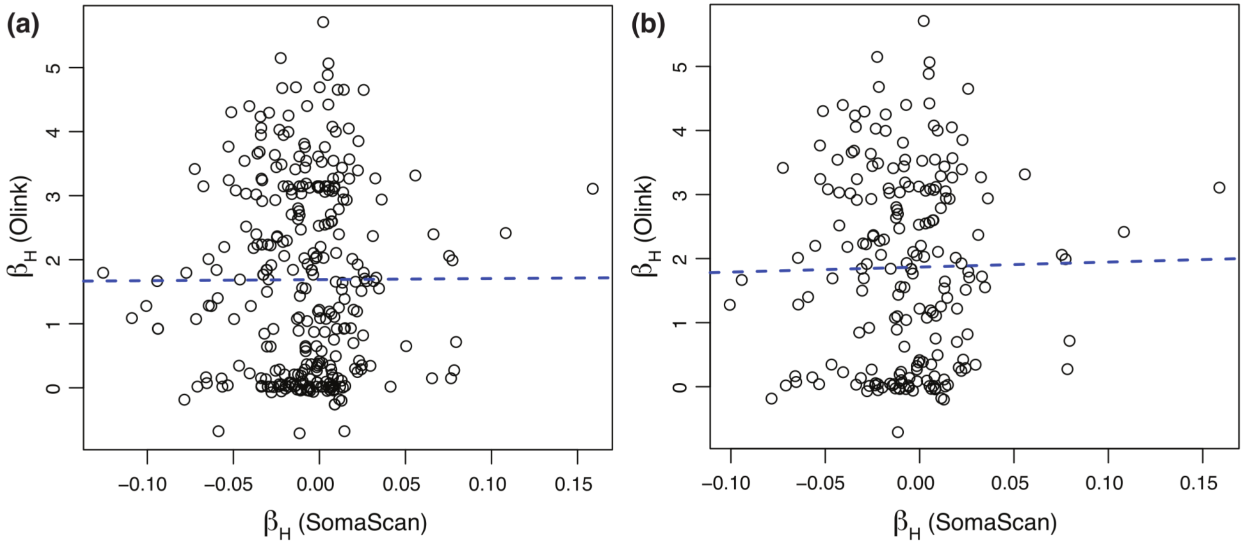


**Supplementary Figure 2. Comparison of hemolysis effects between SomaScan and Olink probes mapped to shared RBC proteins. (a)** Using 305 SomaScan / Olink probe pairs, the correlation is r = 0.004. **(b)** Using a subset of 199 SomaScan / Olink probe pairs (after excluding UniProt IDs that map to more than one SOMAmer or Olink probe), the correlation is r = 0.017. Linear fits are shown as blue dashed lines.
