## Supplementary Figure 3 for "Effects of in vitro hemolysis and repeated freeze-thaw cycles in protein abundance quantification using the SomaScan and Olink assays"

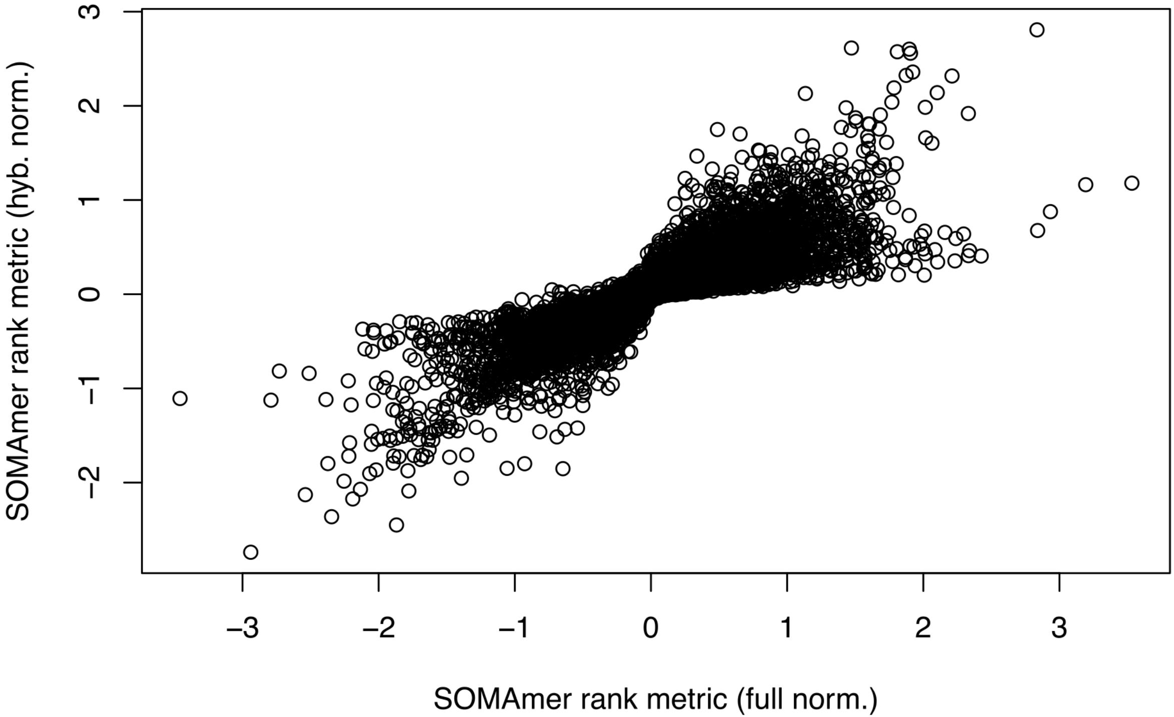


**Supplementary Figure 3. Comparison of SOMAmer rank metrics of hemolysis effects using different levels of data normalization.** SOMAmer rank metrics were defined as -log10(p-value_H)*sign(beta_H). The x-axis shows results obtained using the full normalization (*hybNorm.medNormInt.plateScale.calibrate.anmlQC.qcCheck.anmlSMP*). The y-axis shows results obtained using only the hybridization step (*hybNorm*). The correlation is r=0.867 (p-value<2.2e-16).
