## Supplementary Table 1 for "Effects of in vitro hemolysis and repeated freeze-thaw cycles in protein abundance quantification using the SomaScan and Olink assays"

| OlinkID | UniProt | Assay | Panel | (Intercept) | beta_H | p-value_H |
|---|---|---|---|---|---|---|
| OID21433 | P09105 | HBQ1 | Oncology | 0.66467556 | 3.75914889 | 3.40E-34 |
| OID21078 | Q9NZD4 | AHSP | Neurology | 1.43640565 | 1.21766222 | 8.33E-21 |

| OlinkID | UniProt | Assay | Panel | (Intercept) | beta_H | p-value_H |
|---|---|---|---|---|---|---|
| OID20271 | P07451 | CA3 | Cardiometabolic | 1.360 | 3.416 | 9.76E-33 |
| OID20409 | P00915 | CA1 | Cardiometabolic | 1.953 | 3.031 | 3.86E-54 |
| OID21149 | P00918 | CA2 | Neurology | 2.695 | 1.193 | 1.40E-30 |

**Supplementary Table 1. Hemolysis effects of Olink probes mapped to hemoglobin (top) and carbonic anhydrase (bottom) proteins.**
