## Supplementary Table 2 for "Effects of in vitro hemolysis and repeated freeze-thaw cycles in protein abundance quantification using the SomaScan and Olink assays"

| SeqId | SeqIdVersion | SomaId | Target | UniProt | EntrezGeneID | EntrezGeneSymbol | Dilution | (Intercept) | beta_H | p-value_H |
|---|---|---|---|---|---|---|---|---|---|---|
| 17137-160 | 3 | SL005328 | Beta-globin | P68871 | 3043 | HBB | 5.00E-01 | 3.716 | -1.04E-01 | 0.495 |
| 18198-51 | 3 | SL018342 | HBAT | P09105 | 3049 | HBQ1 | 2.00E+01 | 2.573 | -8.40E-03 | 0.905 |
| 19774-8 | 3 | SL009289 | HBG2 | P69892 | 3048 | HBG2 | 5.00E-01 | 4.334 | -1.16E-01 | 0.359 |
| 26273-24 | 4 | SL017970 | HBAZ | P02008 | 3050 | HBZ | 5.00E-01 | 3.804 | -1.45E-02 | 0.899 |
| 32984-3 | 3 | SL008874 | HBG1 | P69891 | 3047 | HBG1 | 2.00E+01 | 3.920 | -1.31E-01 | 0.444 |
| 4915-64 | 2 | SL000836 | Hemoglobin | P68871\|P69905 | 3043\|3039 | HBB\|HBA1 | 5.00E-01 | 3.762 | -1.49E-01 | 0.412 |
| 6919-3 | 3 | SL017970 | HBAZ | P02008 | 3050 | HBZ | 5.00E-01 | 3.433 | 1.60E-02 | 0.872 |
| 6992-67 | 3 | SL008001 | HBD | P02042 | 3045 | HBD | 2.00E+01 | 2.860 | 1.66E-02 | 0.179 |
| 7965-25 | 3 | SL018342 | HBAT | P09105 | 3049 | HBQ1 | 5.00E-01 | 3.425 | 3.17E-03 | 0.912 |
| 9025-5 | 3 | SL010892 | AHSP | Q9NZD4 | 51327 | AHSP | 2.00E+01 | 3.082 | -6.05E-05 | 0.997 |

| SeqId | SeqIdVersion | SomaId | Target | UniProt | EntrezGeneID | EntrezGeneSymbol | Dilution | (Intercept) | beta_H | p-value_H |
|---|---|---|---|---|---|---|---|---|---|---|
| 3799-11 | 2 | SL004867 | Carbonic anhydrase III | P07451 | 761 | CA3 | 5.00E-01 | 3.527 | -7.24E-02 | 0.511 |
| 4969-2 | 1 | SL004866 | Carbonic anhydrase I | P00915 | 759 | CA1 | 5.00E-01 | 4.645 | -4.26E-02 | 0.502 |
| 4970-50 | 3 | SL000339 | carbonic anhydrase II | P00918 | 760 | CA2 | 2.00E+01 | 2.679 | -7.94E-04 | 0.940 |
| 4970-55 | 1 | SL000339 | carbonic anhydrase II | P00918 | 760 | CA2 | 2.00E+01 | 2.385 | 2.18E-02 | 0.027 |

**Supplementary Table 2. Hemolysis effects of SomaScan probes mapped to hemoglobin (top) and carbonic anhydrase (bottom) proteins.**
